# A computational model elucidates the effect of oncogene induced expression alterations on the energy metabolism of neuroblastoma

**DOI:** 10.1101/2025.02.19.639039

**Authors:** Mareike Simon, Uwe Benary, Katharina Baum, Alexander Schramm, Jana Wolf

**Affiliations:** Mathematical Modelling of Cellular Processes, Max-Delbrück-Center for Molecular Medicine, Berlin, Germany; Department of Mathematics and Computer Science, Free University Berlin, Berlin, Germany; Hasso Plattner Institute, Digital Engineering Faculty, University of Potsdam, Potsdam, Germany; Windreich Department of Artificial Intelligence and Human Health & Hasso Plattner Institute for Digital Health at Mount Sinai, Icahn School of Medicine at Mount Sinai, New York City, USA; Department of Medical Oncology, West German Cancer Center, University Hospital Essen, University of Duisburg-Essen, Essen, Germany

## Abstract

An altered energy metabolism is recognized as a hallmark of cancer. Experimental evidence shows that oncogenes play a key role in the reprogramming of metabolism. In neuroblastoma the oncogene MYCN, a main risk factor of poor prognosis, has been demonstrated to lead to expression changes in numerous glycolytic enzymes. Since it is not clear whether all these targets are required and how the overall metabolic response is shaped, we here dissect the effect of MYCN targets on the pathway individually and in combination using a computational modeling approach. We develop a first mathematical model of the energy metabolism in neuroblastoma cells based on our published experimental data. The analysis shows that MYCN overexpression overall leads to Warburg-like flux alterations with characteristic changes for the individual MYCN targets highlighting that MYCN targets can have opposing and sometimes unexpected effects. Interestingly, not all described MYCN targets contribute to notable flux alterations, at least in glycolysis. Our model moreover predicts a potential bistability of the cellular metabolism with an occurrence of a low-flux state likely representing quiescence. Overall, our study highlights the importance of analysing perturbations such as expression changes in the context of realistic pathways capturing their specific interactions and complex regulations.

## Introduction

The observation that cancer cells harbour an altered energy metabolism dates back to Otto Warburg in the 1920s. According to his hypothesis, cancer cells tend to take up more glucose and convert it to lactate instead of using the more energy efficient pathways of mitochondrial metabolism, even in the presence of sufficient oxygen. This effect was termed aerobic glycolysis or Warburg effect and has been reconsidered only decades after its first description [1]. An altered energy metabolism is now recognized as one of the hallmarks of cancer development [2]. However, it is not entirely understood how the metabolism of cancer cells becomes deregulated and how specific genetic alterations drive metabolic changes. In recent years, this has been the focus of experimental and theoretical research. The impact of oncogenes on the energy metabolism has been intensively studied for various tumour types [3,4]. We here set out to investigate the impact of the oncogene MYCN on metabolic flux alterations in preclinical neuroblastoma models.

Neuroblastoma is one of the most common solid cancers of childhood, accounting for approximately 8% of paediatric cancers and 15% of cancer deaths in children [5]. Neuroblastoma is known for its very heterogeneous prognosis: while the cancer spontaneously regresses in some patients, it presents highly aggressive in others. The survival rate in high-risk cases is still below 50% [6]. One of the major risk factors in neuroblastoma is amplification of the MYCN oncogene that is present in around 20%-25% of the patients [5,6]. MYCN is a transcription factor of the MYC family, which also includes c-MYC and MYCL. While its role in neuroblastoma is known for decades [7], a MYCN amplification was found in 7% of all samples of The Cancer Genome Atlas (TCGA), which represent 33 different tumour types [8]. It was shown experimentally that an amplification of MYCN induces changes in many processes, such as cell proliferation, senescence and inflammation [9]. It was also described that MYCN influences the metabolism in a variety of ways. It alters the expression of glycolytic enzymes [10–12], influences glutamine metabolism [13], β-oxidation [14] and biosynthetic pathways [15]. We and others have shown that MYCN reprograms metabolism on a global scale and enhances glycolysis [16,17]. In an early large scale analysis of genes induced by MYCN four glycolytic genes were identified as MYCN targets: aldolase (ALD), triosephosphate isomerase (TPI), glyceraldehyde 3-phosphate dehydrogenase (GAPDH) and pyruvate kinase (PK) [10]. An upregulation of the lactate transporters (monocarboxylate transporters MCT1 and MCT2) was reported in a later study [11]. In addition, the glucose transporter GLUT1, the enzymes hexokinase (HK), phosphoglycerate kinase (PGK), lactate dehydrogenase (LDH) and pyruvate dehydrogenase kinase (PDK) were identified as upregulated by MYCN [12]. PDK inactivates pyruvate dehydrogenase (PDH), therefore PDH activity is lower if MYCN is overexpressed. Although a plethora of MYCN targets have been described, it is not elucidated why there are numerous MYCN targets within the central energy metabolism, how these interact and to which extent the individual targets contribute to metabolic reprogramming given the complex and multilayered regulation of the energy metabolism. We here address these questions by using a computational model of the energy metabolism and employing experimental data with and without the overexpression of MYCN in a neuroblastoma model [17]. The development of computational models is a well-established method to study metabolic pathways and has been applied to different cell types, including mammalian and yeast cells [18–22]. The investigation of both core and detailed kinetic models has contributed to the elucidation of regulatory principles of the energy metabolism and the impact of intercellular metabolic coupling [18,20,23–25]. Recent modelling approaches also focus on cancer cells, in particular pancreatic [26], cervical [27] and general cancer properties [28]. The model by Shestov et al. [28] motivated the investigation of targeting GAPDH in cells employing aerobic glycolysis [29]. While these cancer specific models are available, to our knowledge, no neuroblastoma-specific model exists. Therefore, this paper focuses on the development of a model of the energy metabolism in neuroblastoma cells based on our experimental data published previously to investigate the effect of MYCN induced expression alteration. The analysis demonstrates a shift of fluxes to a Warburg-like phenotype, but not all experimentally described MYCN targets are required for that. According to our interaction analysis most target effects act additively and only very few target combinations show antagonistic effects. The observation of bistability in MYCN low as well as MYCN high cells might be attributable to differences between metabolic active and inactive cells.

## Results

### A model of the energy metabolism in neuroblastoma

We here set out to develop a model of the energy metabolism that allows for the integration of cell type-specific metabolomics and extracellular flux data obtained from isogenic neuroblastoma cell lines with and without expression of MYCN [17]. For this purpose, we built a model that describes all glycolytic reactions from glucose uptake to pyruvate production, as well as lactate production, secretion and acetyl-coA production in detail. To reduce complexity, the citric acid cycle (TCA), the respiratory chain and the overall ATP consumption are represented by lumped reactions. A scheme of the model is given in Figure 1a. The model incorporates the uptake of the external metabolites glucose and oxygen as well as an exchange of lactate with the medium. The overall model structure is grouped in five parts, marked by coloured vertical bars in Figure 1a: upper glycolysis (dark green), lower glycolysis (light green), lactate production and exchange (blue), ATP consumption (red) and mitochondrial metabolism (purple). It incorporates well-described pathway regulations: HK is inhibited by its product Glc6P, phosphofructokinase (PFK) is regulated by the energy level of the cell, here modelled as inhibition by ATP, and PK is activated by Fru1,6-BP (All metabolite abbreviations are explained in the legend of figure 1). Overall, the model incorporates 18 reactions and 19 metabolites. There are three conserved moieties; a conservation relation for ATP and ADP, one for NAD and NADH, and a third conservation capturing the total phosphate level and therefore including phosphate, ATP and all glycolytic intermediates from Glc6P to PEP. Details of the model are given in the Supplementary Material.

**Figure 1.**
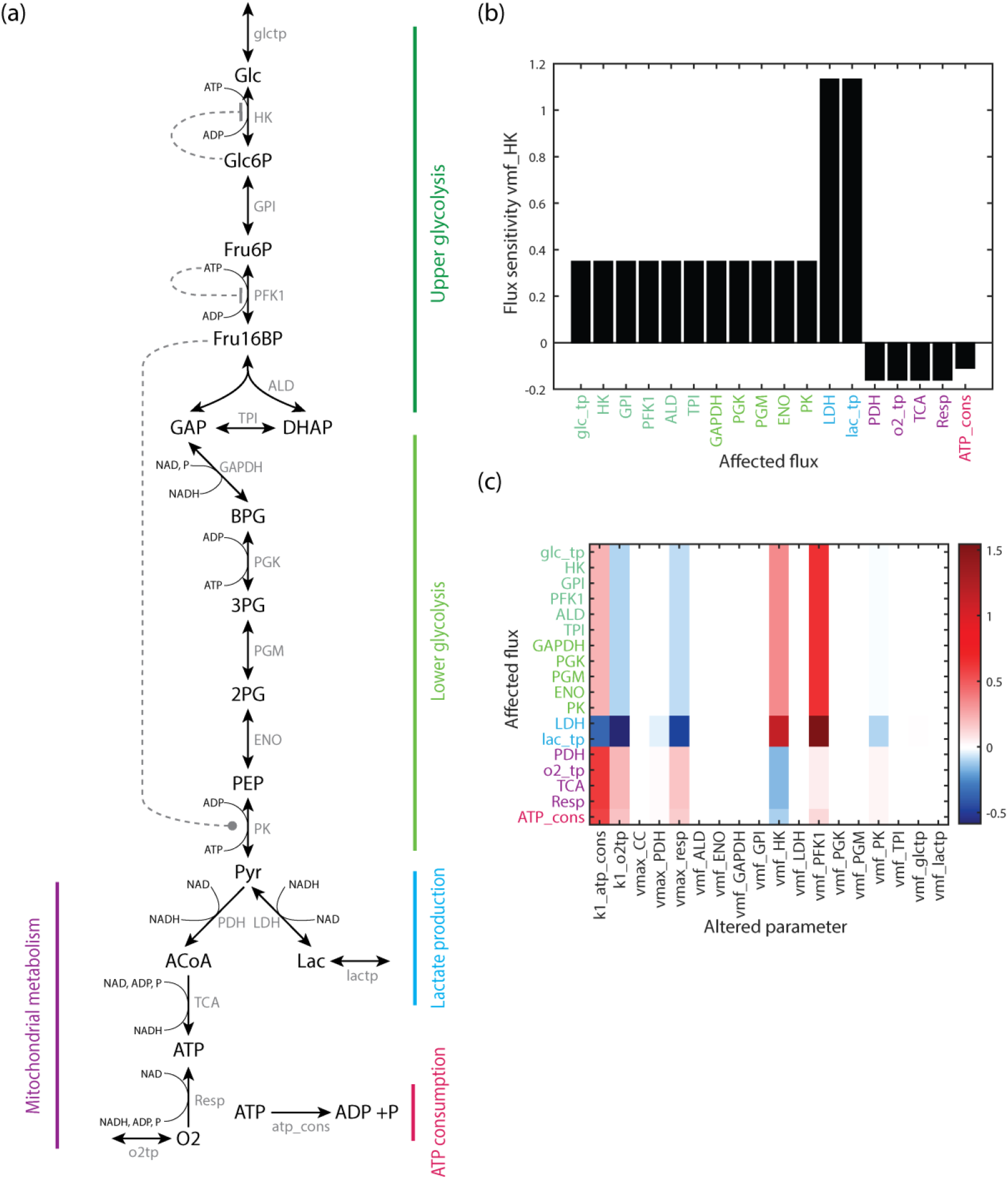
Model structure and sensitivity of fluxes. (a) Structure of the computational model of the energy metabolism. The model contains the following metabolites: Glc glucose, Glc6P glucose 6-phosphate, Fru6P fructose 6-phosphate, Fru16BP fructose 1,6-bisphosphate, GAP glyceraldehyde 3-phosphate, DHAP dihydroxyacetone phosphate, BPG 1,3-bisphosphoglycerate, 3PG 3-phosphoglycerate, 2PG 2-phosphoglycerate, PEP phosphoenolpyruvate, Pyr pyruvate, Lac lactate, ACoA acetyl-coenzyme A, P Phosphate, O2 O_2_, as well as ATP, ADP, NAD and NADH. The external metabolites glucose glc_ext, lactate lac_ext, and oxygen O2_ext are not shown. The included reactions are: glctp glucose transport, HK hexokinase, GPI glucose 6-phosphate isomerase, PFK phosphofructokinase, ALD aldolase, TPI triosephosphate isomerase, GAPDH glyceraldehyde 3-phosphate dehydrogenase, PGK phosphoglycerate kinase, PGM phosphoglycerate mutase, ENO enolase, PK pyruvate kinase, LDH lactate dehydrogenase, lactp lactate transporter, PDH, pyruvate dehydrogenase, TCA (or CC) tricarboxylic acid cycle, Resp Respiration, o2tp oxygen transport, atp_cons ATP consumption. Arrows represent reactions or transport processes, dashed lines regulations. The overall pathway can be divided in five parts, which are marked by coloured vertical bars: upper glycolysis (dark green), lower glycolysis (light green), lactate production and exchange (blue), mitochondrial metabolism (purple) and ATP consumption (magenta). (b) Normalized flux sensitivity coefficients for changes in the maximal velocity of HKat the reference state of the baseline model. Flux sensitivities are equal within the pathway parts shown in panel a: upper and lower glycolysis, lactate production, mitochondrial metabolism and ATP consumption. (c) Sensitivities of all model fluxes to the kinetic parameters. Given are normalized sensitivity coefficients of the model fluxes (y-axis, labeled by enzyme name of the reaction) for changes in parameters (x-axis, all maximal velocities and rate constants). Red colours show positive sensitivity coefficients, blue negative sensitivity coefficients.

All but two reactions follow Michaelis-Menten type kinetics, and these concern oxygen transport and overall ATP consumption, which are represented by mass action kinetics. While most reactions are modelled as reversible reactions, the mitochondrial reactions and ATP consumption reactions are represented as irreversible processes. The detailed kinetic descriptions are given in the supplementary material. The kinetic parameters are estimated from neuroblastoma cell line based metabolomics data and extracellular flux measurements, supplemented by parameter values from literature. Our metabolomics data capture the metabolite levels in SH-EP neuroblastoma cells with Tet-inducible MYCN expression, which were grown under three different glucose concentrations [17]. In addition, the extracellular acidification rate (ECAR), characterizing the lactate release of the cells, and the oxygen consumption rate (OCR) were measured for these conditions using a Seahorse analyser. Since maximal velocities are commonly assumed to be cell type specific, we used the neuroblastoma specific metabolomics and flux data to estimate these values by parameter fitting. Details on the parameter assignment and fitting procedure are given in Materials and Methods.

The parameter set of the best model fit is given in Supplement Table S.1. The model utilizing that parameter set will be referred to as baseline model, the fitted steady state as reference state. A comparison of its metabolite concentrations with respective literature values shows that all values are in the expected range, see Figure S.1. The values of steady state metabolite concentrations and fluxes are given in Supplemental Tables S.2 and S.3.

### Sensitivity analysis of the baseline model shows strong impact of kinases, respiration and ATP consumption

To find out how strongly a change in one reaction impacts all other reactions, we analysed the sensitivity of the steady state fluxes towards parameter changes. Figure 1b shows the effect of a parameter change in the maximal velocity of HK. It can be observed that groups of fluxes respond similarly, that holds for processes of upper and lower part of glycolysis (dark and light green in Fig. 1a), for lactate production and exchange (blue bar in Fig. 1a), and for the mitochondrial metabolism (purple bar in Fig. 1a), respectively. Steady state fluxes within these groups show the same sensitivity towards parameter changes, which result from stoichiometric constraints within the pathway leading to linear dependencies of fluxes. Therefore, four exemplary fluxes are sufficient to characterize the response of the system. We chose glucose uptake, lactate release, oxygen uptake and ATP consumption (red bar in Fig. 1a) for that purpose.

Next, we analysed how strongly the parameters of all reactions influence the steady state fluxes of the system. Figure 1c shows the impact of parameters, in particular the maximal velocities and rate constants, on all steady state fluxes. This demonstrates that the parameters of six processes have the strongest impact, that is of ATP consumption (k1_atp_cons), oxygen transport (k1_o2tp), respiration (vmax_resp) and the kinase mediated reactions (vmf_HK, vmf_PFK1, vmf_PK). The other parameters have a very small or no impact (Fig. 1c, white colour). Notably, the maximal velocities of the kinases HK and PFK have a strong impact on the flux distribution. While an increase in the activity of both kinases leads to increased fluxes through glycolysis and lactate production (red colour in Fig. 1c), one can observe a decrease in the mitochondrial metabolism and ATP consumption for HK (blue colour), while an increase in PFK activity leads to an increased flux through these reactions. The sensitivity coefficients of the maximal velocity of the PK reaction are generally smaller and show a decrease in glycolytic and lactate fluxes combined with an increase in the fluxes of mitochondrial reactions and ATP consumption. This pattern can qualitatively also be observed for the oxygen transport and the respiration process. The ATP consumption reaction has a strong effect on the steady state flux distribution, with a negative effect on the lactate related reactions and a positive effect on all other reactions. Overall, this analysis shows that the maximal velocities of the kinases, respiration and ATP consumption have specific and strong impacts on the steady state flux distribution.

### The baseline model shows bistability with an additional, low-flux state

Since the occurrence of bistability was observed in glycolysis of yeast [30] and mammalian cells [31], we investigated the co-existence of steady states in our model. Because the complexity of the derived model does not allow for a full bifurcation analysis, we investigated the existence of stable attractors by a simulation strategy: for that purpose, parameters are altered slightly and stable steady states are identified by time course simulations. Details of the approach are given in the Material and Method Section. There is a wide parameter region where two stable steady states coexist, as exemplified for the maximal velocities of HK (Fig. 2a) PFK and PDH (Suppl. Fig S.2). In addition to the steady state that fits the experimental data best (reference state, upper branch in Fig. 2a), a second steady state exists, which is characterized by a very low glucose uptake rate (lower branch in Fig. 2a). The parameter value of the reference state (Fig. 2a, broken vertical line) lies within the parameter region of two coexisting stable states, whereas for high vmf_HK values only the steady state with a low glucose uptake rate exists.

**Figure 2.**
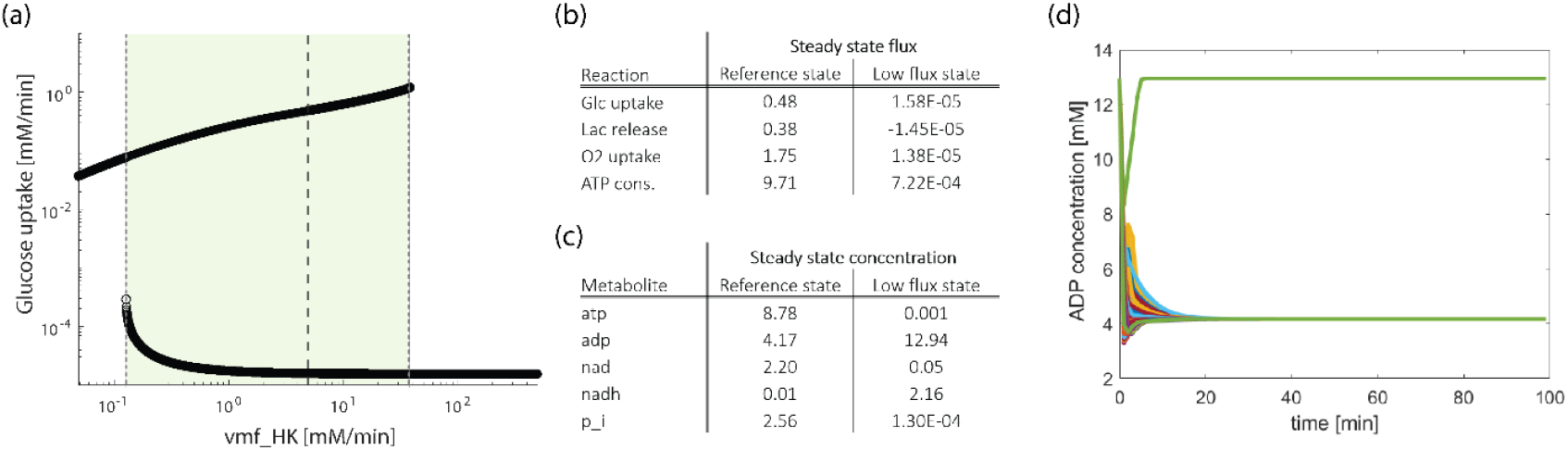
Coexistence of two stable steady states. (a) Stable steady state values in dependence of one parameter, that is the maximal velocity of the hexokinase reaction. The dashed line indicates the fitted value of the parameter vmf_HK. All other parameters are at their reference values (Supplement Table S.1). The shaded area confined by dotted lines marks the parameter region of coexisting steady states. (b) Fluxes and (c) concentrations in the two steady states for the reference parameter set. (d) Time course simulations of the ADP concentration at the reference parameter set for different initial concentrations.

We next investigated the fluxes in this alternative state (Fig. 2b). All of them, including glucose and O_2_ uptake as well as ATP consumption were considerably low, in line with a quiescent state of the cells. We therefore designated this as “low flux state”. It is also characterized by a lactate uptake instead of a secretion, indicated by a negative value of the lactate release flux and suggesting that lactate fuels the mitochondrial metabolism. An investigation of the steady state concentrations under this condition demonstrated that almost the entire ATP-ADP pool shifted to ADP with only minimal levels of ATP (Fig. 2c). For the redox pool a high concentration in NADH can be observed, while NAD levels are very low, which is in strong contrast to the reference state. Also, the phosphate concentration in the low flux state is reduced. This shows that in all conserved moieties rather extreme ratios are reached. Furthermore, concentrations within the modules of the upper glycolysis are increased, and correspondingly very low for the lower glycolysis (Suppl. Table S.2). Our results suggest GAPDH to serve as a bottleneck, as two of its substrates (NAD, phosphate) are present in very low concentrations.

In order to investigate the attractor basins of the two stable steady states, the model was simulated from various initial concentrations. We randomly assigned 2000 initial concentrations obeying the constraints of the conserved moieties. In these simulations, all trajectories reach the reference state. In a second set of simulations, initial concentrations of metabolites were assigned randomly but ATP and phosphate concentrations were restricted to very low initial values (0.001 mM). In this case, simulations from 2000 initial conditions show that both stable steady states can be reached (Fig. 2d, exemplified by the high and low ADP concentration of the two states). This verifies the existence of the two stable steady states and highlights that the low flux steady state has a rather restricted basin of attraction.

### Implementation of MYCN target effects

The establishment and analysis of the baseline model, which represents a low MYCN state, serves as a basis to investigate the effects of MYCN overexpression. As outlined in the introduction there is a high number of MYCN targets described in the literature. For our model the transporters of glucose and lactate as well as the enzymes HK, ALD, TPI, GAPDH, PGK, PK and LDH are relevant. Moreover, PDK is a described

MYCN target. Since PDK inactivates PDH, PDH activity is lowered if MYCN is over-expressed. These MYCN targets are visualized in Figure 3a, illustrating their broad distribution over the pathway.

**Figure 3.**
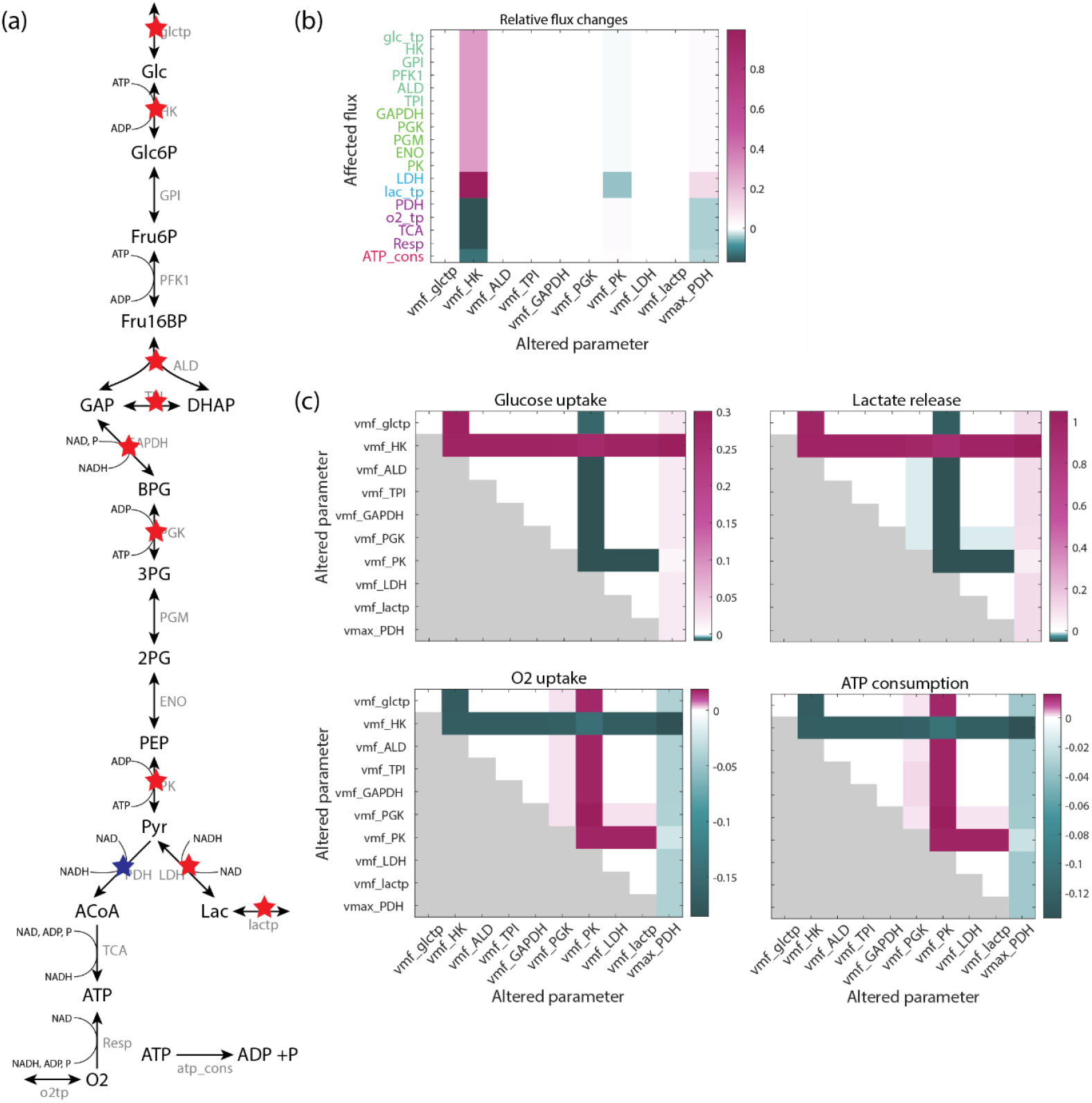
Effect of MYCN targets on flux distribution. (a) Visualization of literature derived MYCN targets in the pathway model. Red and blue stars mark the enzymes with a described expression increase and decrease in MYCN dependence, respectively. (b) Effect of individual MYCN targets on the flux distribution for a MYCN factor of 2 (the PDH activity is decreased, all other enzyme activities are increased); white corresponds to an unchanged flux; purple and green show increases and decreases in the fluxes, respectively.(c) Effect of pairwise MYCN targets on the flux distribution for a MYCN factor of 2. The diagonal elements of each subfigure depict the individual effects for comparison. Only half of each subfigure is filled due to symmetry.

In order to implement the MYCN effects in the model, maximal velocities of the respective reactions are changed. This is motivated by the fact that maximal velocities are defined as *v*_*max*_ = *k*_*cat*_ *·* [*E*], with enzyme concentration [*E*], taking a linear relation between the enzyme concentration and the maximal velocity for each reaction into account. Since MYCN induces the expression of the described enzymes, the maximal velocities were increased except for PDH, which was reduced due to described effects of the increased expression of PDK. Therefore, the maximal velocities are multiplied by a ‘MYCN factor’, for the maximal velocity of PDH a reciprocal factor was applied. So far, there are no data to explicitly derive the size of these factors for each reaction directly from experiments. In earlier studies including our own [17], changes in the specific activity of some of the enzymes had been observed to be rather small. On the other hand, analyses of the gene expression levels mostly showed stronger induction [10–12]. Therefore, we here mostly investigated MYCN factors in the range [1,2].

### Effects of individual and paired MYCN targets

In the following we addressed the question how the individual MYCN targets affect the pathway behaviour and what their combined effect is. The impact of individual MYCN targets on the model behaviour was investigated by applying a MYCN factor that represents MYCN-induced changes in protein expression and subsequent effects on the maximal velocity of the respective protein. Next, the resulting effects on the steady state flux distribution were analysed. For example, a MYCN factor of 2 induces strong changes in flux distribution for only three out of the ten MYCN targets, HK, PK and PDH with strongest changes in the activity of HK (Fig. 3b). This leads to an increase in glycolytic and lactate production fluxes (purple bars) and a decreased flux through mitochondrial metabolism and ATP consumption (green bars). Changes in the activity of PK also shows an effect, which is opposite to that of HK, thus causing a decrease in glycolysis and lactate production (green bars) and a slight increase in mitochondrial metabolism and ATP consumption (light purple bars). This suggests that these are partly ATP/ADP mediated effects, since ATP is a substrate for the HK reaction, while ADP is a substrate of PK. A third enzyme with a notable effect on the flux distribution is PDH. A reduction in the rate of this enzyme, which catalyses the first reaction in the mitochondrial module in our model, leads to a reduced flux through mitochondrial metabolism (green bar). Moreover, it slightly increases the flux through glycolysis and lactate production (purple bars), and decreases ATP consumption (light green bar). Applying a factor for the other MYCN targets do not lead to notable flux alterations (white bars in Fig. 3b for a MYCN factor of 2). The result is similar for other values of the MYCN factors in the range [1,2]. Overall, this supports the sensitivity analysis of the baseline model performed for very small parameter perturbations which also indicated that changes in the maximal velocities (vmax) of ALD, TPI, GAPDH, PGK and LDH as well as of the glucose and lactate transport hardly induce any flux changes, compare Figure 1c.

For the analysis of paired MYCN targets we investigated the effect on the earlier defined representative fluxes glucose uptake, lactate release, oxygen uptake and ATP consumption. In the analysis two enzyme rates were simultaneously altered by the same factor and the changes in the steady state fluxes were observed. The results were visualized in individual subplots of Figure 3c, where in each case the diagonal elements represent the impact of individual alterations, allowing for an easier comparison with Figure 3b. Due to the symmetry of the analyses, only half of each subfigure is filled.

The subplots of Figure 3c show that the three MYCN targets HK, PK and PDH which dominate when individually changed, also shape the response of the system for combined perturbations. The enzymes and transporters that played no substantial role individually mostly also have no strong contribution in the combined cases (white boxes, Fig. 3c). An exception is PGK which shows a small effect on lactate release, oxygen uptake and ATP consumption fluxes that was not observable for individual perturbations, mostly due to the scaling.

The effect of combined perturbations is often dominated by one of the perturbations, e.g. for the pair HK and ALD. However, in some cases a modulation of the effect of both perturbations can be observed, e.g. for PDH and PK, or HK and PK. This raises the question whether the effect of MYCN targets is always just additive. In order to study different possibilities of target interaction, we used an algorithm that classifies interactions as one of three types: additive, if the flux changes are given by the sum of the two individual changes; synergistic, if the change is higher than the sum of the two; and antagonistic if the change is smaller than the sum of the two changes. Details of the approach, that was adapted from Piggott et al. [32] are given in the Material and Method Section. Supplementary Figure S.3 shows the results of this interaction classification for a MYCN factor of 2. We find the majority of changes for MYCN targets pairs to be additive. Interestingly, in two cases antagonistic interactions were detected. These are HK changes paired with changes in PK and PDH enzymes. However, we did not detect any synergistic effects between MYCN targets. Similar pictures arise for MYCN factors of 1.5 and 5. Overall, this shows that the effect of MYCN targets is not always additive, but also antagonism can arise demonstrating that the impact of two targets can have counteractive effects on pathway fluxes beyond their individual contributions.

### The MYCN high model shows Warburg-like changes in steady state fluxes

In the case of MYCN overexpression likely all MYCN targets are simultaneous influenced. This was implemented by applying a MYCN factor to all targets simultaneously, the resulting model will be referred to as MYCN high model. In a first scenario we used the same factor for all MYCN targets and compared the model behaviour to that of the baseline model. Simulations of the MYCN high model were performed for MYCN factors of 1.5 and 2.

The steady state fluxes and concentrations of the corresponding MYCN high model are listed in Supplemental Tables S.3 and S.4. The high MYCN model shows an increase in the glucose uptake and lactate release fluxes with increasing MYCN factor (Table S.3). In contrast, oxygen uptake and ATP consumption decrease with increasing MYCN factor, demonstrating that the MYCN high models show a Warburg-like phenotype. The strongest relative change can be observed in lactate release compared to moderate changes in the other fluxes.

We next investigate the flux sensitivities of the MYCN high model. The result for a MYCN factor of 2 is given in Figure 4a. It shows that the MYCN high model is sensitive to changes in the maximal velocities of the kinases HK, PFK, PK and atp_cons that were also prominent in the baseline model. For these parameters the sensitivity of glycolysis and lactate production are lower, while that of the mitochondrial metabolism is increased compared to the baseline model. A striking observation is the strongly reduced sensitivity for k1_o2tp in the MYCN high model indicating that this model becomes more independent from oxygen. In addition, the sensitivity towards the maximal velocity of PDH increases in the MYCN high model. The other maximal velocities have no significant impact on the fluxes, shown by their very low sensitivity coefficients (white colour in Fig. 4a), which is similar to their impact in the baseline model. Overall, the baseline and the high MYCN model are mostly sensitive towards the same parameters, with one prominent exception, that is the sensitivity towards the oxygen uptake rate of the cell.

**Figure 4.**
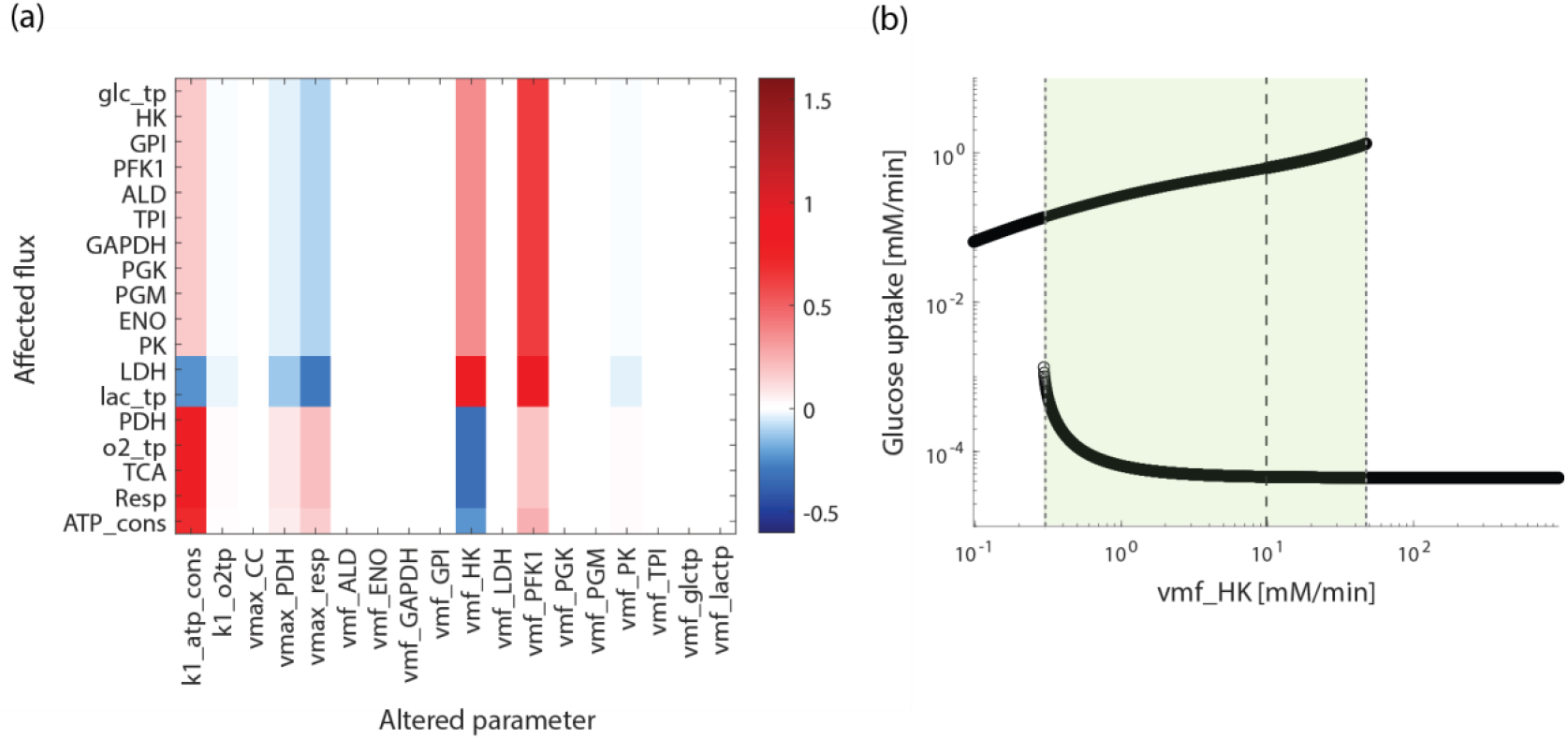
Flux sensitivities and bistability in the MYCN high model. (a) Normalized flux sensitivity coefficients for all maximal velocities and rate constants of the MYCN high model; MYCN factor 2. (b) Stable steady states in dependence of the maximal velocity of the hexokinase reaction for a MYCN factor of 2. The dashed line indicates the starting parameter value, the dotted lines mark the parameter values where one of the steady states becomes unstable.

It is an interesting question whether the MYCN high model can also exhibit multi-stationarity. Figure 4b shows the result of the corresponding analysis, demonstrating that the MYCN high model has an additional steady state characterised by a very low glucose uptake rate (Fig. 4b). Overall, the bifurcation-like diagram is very similar to that of the baseline model with two stable steady states coexisting for a broad parameter range, compare Figure 2a. A detailed comparison of the bistabilities reveals quantitative differences between the baseline and MYCN high model: it shows that (i) the glucose uptake flux in the alternative steady state is slightly increased in the MYCN high model and (ii) the parameter range of vmf_HK for co-existing steady states is slightly smaller for the MYCN high model. These analyses together highlight that the general behaviour between the baseline and MYCN high model is qualitatively similar, but quantitative changes occur.

### Individually assigned MYCN factors predict range of characteristic flux changes

While we so far analysed cases where all MYCN targets are altered by the same factor we now study how the flux distribution changes when each of the MYCN targets is altered by an individual factor. To that end, we assigned random values for the MYCN factors to each MYCN target. The values are drawn from a uniform distribution of a given range and steady state flux simulations are performed for 100.000 assignments of the factors. Figure 5a shows the results for MYCN factors of the range 1 to 2. This demonstrates for nearly all simulations an increase in glucose uptake and lactate release accompanied by a decrease in oxygen uptake and ATP consumption rates compared to the baseline model (red broken line in Fig. 5a) is observed. Only in very few cases a decrease in glucose uptake and lactate release occurs, increases in oxygen uptake and ATP consumption are slightly more common.

**Figure 5.**
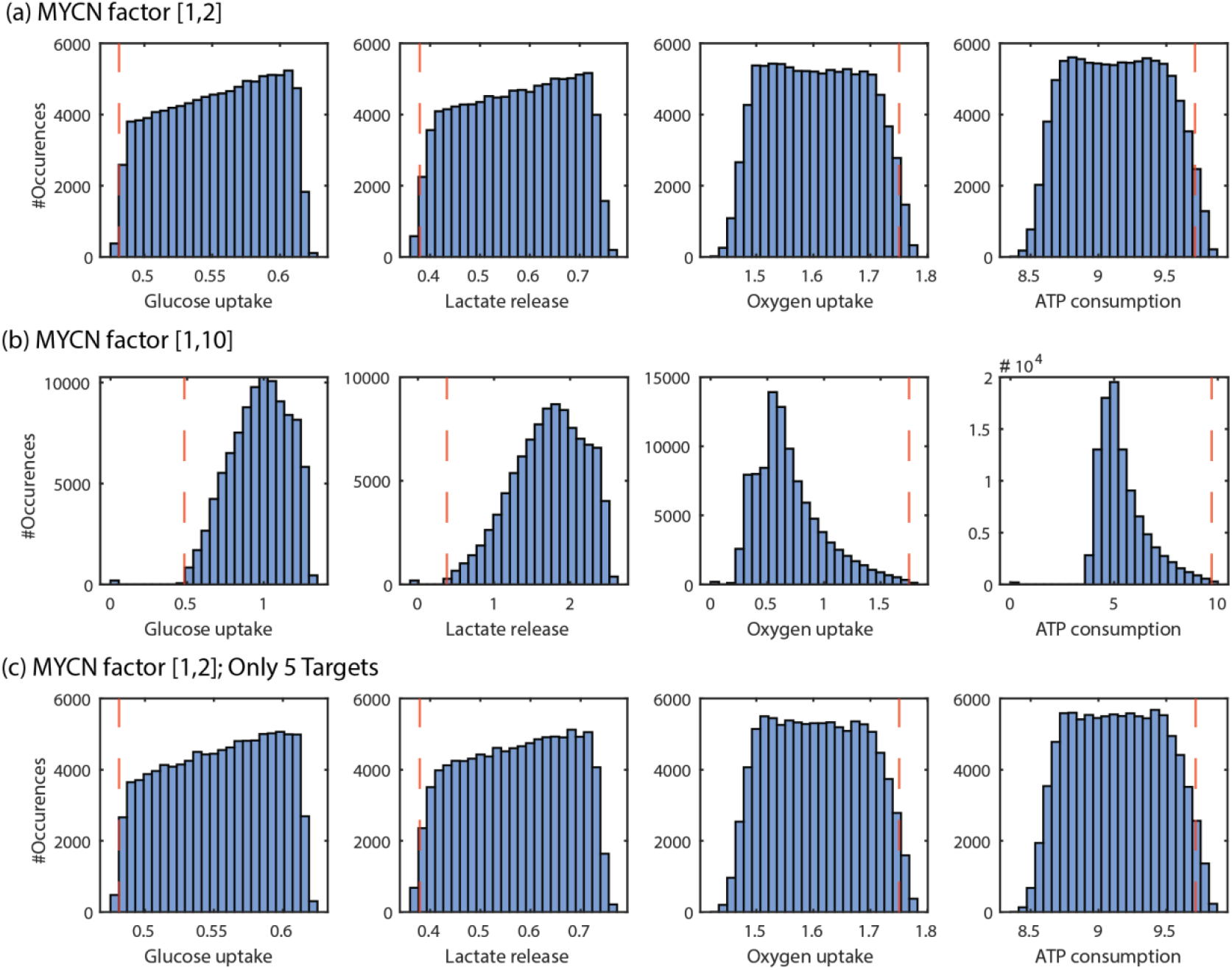
Characteristic steady state fluxes for variable MYCN factors. Flux distribution for randomly assigned factors of each MYCN target. Values assigned from a pre-defined range: (a) [1,2], (b) [1,10]. The model is simulated 100,000 times for individually assigned random factors for the MYCN targets. For PDH the activity is decreased, all others are increased. The red dashed line indicates the corresponding flux in the fitted baseline model. All fluxes in [mM/min]. (c) Only five of the described MYCN targets, that is glctp, HK, PGK, PK and PDH, are altered by a randomly chosen individual value from the range [1,2].

In order to study the flux changes for higher MYCN factors, that is stronger MYCN effects, the analysis was repeated for MYCN factors from the range of 1 to 10 (Fig. 5b). In that case the flux distribution is shifted towards even higher values for glucose uptake and lactate release as well as lower values for oxygen uptake and ATP consumption. Interestingly, here some simulations lead to the alternative steady state representing the low flux state. In these cases, all four fluxes are close to zero. Overall, the analyses highlight that MYCN over-expression leads to an increased flux through glycolysis and an increased lactate production with a decreased activity of the mitochondrial metabolism. This represents a Warburg-like phenotype of glycolysis. Stronger MYCN effects can also lead to the alternative state.

### Not all MYCN targets are required for the induction of characteristic flux changes

Since the earlier analysis showed that not all described MYCN targets have a strong effect on the flux state (see Fig. 4a), we investigated the steady state fluxes of the MYCN high model via a ‘leave one out’ analysis, in which all targets except one are altered.

The results of this analysis are given in Table S.5 in the Supplement for a MYCN factor of 2. It shows that the MYCN targets ALD, TPI, GAPDH, LDH and lactp can be left out without a noticeable change in the flux distribution of the model. Only leaving out one of the MYCN targets glctp, HK, PGK, PK and PDH change the characteristic fluxes (highlighted in grey in Supp. Table S.5). The neglectable impact of ALD, TPI, GAPDH, LDH and lactp was confirmed by a simulation where these targets were left out simultaneously. Indeed, leaving out all five does not alter the steady state flux distribution significantly, see Table S.5. In order to verify the result that glctp, HK, PGK, PK and PDH are sufficient for an altered flux distribution in the MYCN high model, the analysis for individualised alterations of the MYCN factors performed in Figure 5a was repeated but only these five targets, altered by a random factor between 1 and 2. The resulting flux distribution, shown in Figure 5c, is very similar to that in Figure 5a where all MYCN factors have been modified. Taken together these analyses demonstrates that not all MYCN induced alterations are required to achieve Warburg-like flux changes.

## Discussion

The oncogene MYCN is well known for its wide range of cellular targets and has been shown to be critically involved in reshaping cellular metabolism. The number of literature-described MYCN targets within the core energy metabolism is surprisingly high, raising questions how the multiple targets interact and jointly shape the MYCN effect. By establishing a kinetic model for the energy metabolism in neuroblastoma cell lines and analysing all described MYCN targets we demonstrated that while MYCN expression leads overall to a Warburg-like shift in the fluxes, the impact of individual targets differs significantly (Fig. 3). We found that several targets have minimal impact on the fluxes, while others change the fluxes in an enzyme-specific way. The strongest flux changes can be observed for the MYCN targets HK and PK as well as PDH which is an indirect MYCN target, regulated via pyruvate dehydrogenase kinase. An interaction analysis of the targets demonstrates that their effects are mostly additive, but antagonistic effects can be observed for the pairs HK-PK and HK-PDH (Fig. S3). We did not observe any synergistic effect between MYCN targets on the fluxes of the energy metabolism.

In simulations where all MYCN targets are simultaneously modified by individual MYCN factors a wide range of flux changes occurs (Fig. 5a). While in the majority of cases the Warburg-typical increase in glucose uptake and lactate release combined with a decrease in oxygen uptake and ATP consumption occur, there are also cases where the direction of flux changes varies. This might reflect variability of conditions or cell lines, as for instance reported by Oliynik et al. [16] describing increased ECAR and OCR values in response to MYCN over-expression in a different neuroblastoma cell line. In this context is would be interesting to further explore the impact of intra-or inter-tumour heterogeneity [33,34] by introducing variability in the model parametrization. To account for this, an experimental characterisation of multiple neuroblastoma cell lines under comparable conditions will be required. Furthermore, one could take into account spatial variability within tumours that involves availability of nutrients or oxygen as it has been reported that MYCN directly impacts on the cellular oxygen sensing system by interacting with HIF-1 in hypoxic neuroblastoma cells [12].

An interesting observation of our computational model is that both the baseline and the MYCN high model show bistability. The occurring additional stable steady state is characterized by very low fluxes, characterizing this state as likely quiescent state of cells. These could be linked to metabolically inactive, dormant or senescent cells. Simulations showed the basin of attraction of the alternative state in our model to be rather small, but low initial ATP and phosphate levels can shift the system towards these states. It will be interesting to explore the bistability, both experimentally and theoretically, for different scenarios, e.g. transient signals or media compositions, to predict under which conditions MYCN high cells can be targeted to shift them to quiescent states. While the concept of coexisting metabolic active and inactive states has been computationally and experimentally studied in detail in yeast [30] it has been only very recently shown for a case of targeting hepatocellular carcinoma [35]. In-depth analysis in yeast showed that it arises from an imbalance of ATP producing and consuming reactions in the glycolytic pathway and represents a metabolic variability with nongenetic origin. Moreover, the possibility of multistability in glycolysis has been experimentally and computationally shown to be possible due to different isoform compositions of glycolytic enzymes [31].

We used our established model to compare the flux sensitivities with and without over-expression of the MYCN oncogene. Despite the overall shift in fluxes, we found the sensitivities to differ only slightly in the baseline and high MYCN models. The only strong deviation is a substantially decreased sensitivity of the MYCN high model towards the oxygen transport. This shows that in our model MYCN over-expression does hardly create distinct vulnerabilities towards targeted therapies within glycolysis despite its high number of MYCN targets. Our analysis moreover demonstrated that not all targets are required to remodel the flux distribution. The leave-one-out analysis showed that a number of MYCN targets has minimal effect of the central fluxes (Fig 5c, Table S.5). One can contemplate that these targets may be involved in the regulation of side branches of glycolysis or connected pathways, e.g., serine-glycine-one-carbon pathway, serine synthesis or glutathione metabolism, which have been reported to be impacted by MYCN over-expression in experiments [17,36,37]. To study the impact of MYCN in larger metabolic networks, comprehensive metabolic models including side branches and biosynthetic processes are required. However, most kinetic cancer models so far focus on the central energy metabolism, some with extension to glycogen or glutamine metabolism [26,28,38,39]. To foster the understanding of cancer metabolism hybrid modelling combining detailed kinetic models with genome scale models might be an interesting future approach. Our study lays the foundation for a better understanding of MYC-driven metabolic changes that might help to identify critical nodes in tumour metabolism that could be exploited in future therapeutic approaches.

## Materials and Methods

### Processing and incorporation of experimental data of Neuroblastoma cell lines

Our model is based on an experimental characterization of SHEP-TR_MYCN neuroblastoma cells by mass spectrometry-based metabolomics measurements and flux measurements by Seahorse technology. The methods previously described in detail [17] provide metabolite levels, extracellular acidification rates and oxygen consumption rates under low and high MYCN expression as well as three different glucose concentrations in the medium. The model fitting is based on the data for low MYCN expression only. For these conditions metabolite levels were quantified for a total of 499 metabolites in three replicates [17]. The metabolomics data was normalized to cell count and median for scaling. Data for 12 metabolites without imputation of missing values are used for model fitting. Scaling and error parameters are fitted once for each metabolite. From three independent extracellular flux measurement, which quantify the acidification rate of the medium and the oxygen uptake of the cell over time, we aimed to extract steady state values for the lactate secretion and oxygen uptake. Therefore, we consistently used the third time point of the experiments, since at the first two timepoints a steady state is not yet reached, later time points do not represent unperturbed state in all experiments. The oxygen consumption rate (OCR) was corrected for non-mitochondrial uptake by subtracting the value of last time point of the experiment after Rotenone treatment. The extracellular acidification rate (ECAR) was correlated to lactate uptake as described in literature [40]. The buffering power of all three media was determined by linear fits. The fitting of the parameter maxH/O2 was restricted to the range 0.65 to 1, based on the range given in literature [40]. The pH value was set to the mean pH value of the experiment. For each experiment individual scaling parameter which were the same for OCR/ECAR are fitted and can be interpreted as a conversion from pmol/min to mM/min, but the same error parameters for all experiments were fitted. Since no glucose uptake experiments were available, a literature derived value is used [41], the maximal velocities could be scaled to represent different absolute rates. The parameter boundaries for the maximal velocities were on a logscale [-2,4], which corresponds the typical range in similar models. The Michaelis-Menten constants for condensed reactions (TCA cycle, respiratory chain) were also estimated by fitting, since the parameters of condensed reactions cannot be directly derived from databases. For these the boundaries are [-3,1] based on the typical values for this parameter type. Additional kinetic parameter such as k_m_ values were taken from the databases BRENDA [42] and SABIO-RK [43]. For equilibrium constants literature data was acquired [44]

### Parameter fitting

Parameters that could not be derived from literature and databases were fitted using the Data2Dynamics (D2D) Toolbox [45]. The starting values for the Michaelis-Menten constants are drawn from a random distribution between 10^−3^ and 10^1^, the other parameters are fitted between 10^−2^ and 10^4^. The range for initial concentrations of the metabolites are based on literature values [26,46]. The external glucose concentration is known in the experiments, external lactate was set to 0.5 mM, which is comparable to other models and the external oxygen concentration was set to 0.181 mM [47]. For the fitting 25000 initial parameter sets were sampled using latin hypercube sampling and then optimized using the default algorithm LSQNONLIN. The model was fitted to the pre-processed metabolomics data and Seahorse measurements as described in the Method Section ‘Processing and incorporation of experimental data of Neuroblastoma cell lines. In both cases scaling factors and error parameters are estimated. Since the data sets represent steady state conditions, the model is equilibrated towards a steady state for fitting. This is achieved using the arSteadyState function available in the toolbox [45]. In order to reach feasible steady state concentrations, the function SteadyStateBounds is used, which adds a quadratic penalty when the steady state concentration for parameter sets falls outside of defined boundaries. The chosen boundaries for the metabolites are derived from literature [26] and represent typical concentrations in various mammalian cell lines, mostly representing cancer.

### Model simulations

All simulations were performed in MATLAB 2019a, using either the D2D framework or the inbuilt ODE solver ode23s. All model analyses in this paper were done using a medium glucose concentration of 11.1 mM, which is the intermediate concentrations used in the experiments [17]. Model investigations were performed for the best model fit, the analyses of individual and pairwise MYCN factor impacts as well as random changes in the MYCN factor were repeated for the second and third best fit with similar results. The MYCN related analysis was performed for MYCN factors of 1.5 and 2, showing the same trends in the model behaviour.

### Sensitivity coefficients

Sensitivity coefficients s of the readout y, here the steady state flux, and the parameter p are calculated 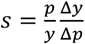. They are calculated for a 1% increase in the parameters.

### Analysis of bistability

Due to the model complexity, direct bifurcation analyses cannot be performed. In order to analyse the possible bistability of the model, an alternative strategy was used: the model was simulated after stepwise alterations of individual parameters. Overall, parameters were changed by a factor of 100 and 1/100. The overall parameter range was divided in 1000 steps on a log scale. For each step, the model is simulated for 100000 mins, to ensure that the systems reached a steady state. The final state of the previous simulation was used as the initial concentration for the simulation.

### Random assignment of fold changes for MYCN targets

For each of the 10 described MYCN targets a random fold change value is drawn from a univariate distribution within the given borders [1,2] or [1,10] (see Fig. 5a, b). For that the inbuilt MATLAB function rand is used. The model is then simulated for each set of parameters with D2D [45]. This is repeated 100000 times resulting in a distribution of fluxes.

### Interaction analysis

For the interaction analysis, we adapted the approach of Piggott et al. [32], which was developed to investigate the interaction between multiple stressors and analyses positive and negative changes. We adapted the method originally applied to systems in ecology, since it considers changes in both directions contrary to many interaction analysis methods classically used for drug interactions. Here we concentrate on three potential interactions types: additive, if the overall change is the sum of the two individual changes; synergistic, if the overall change is higher than the sum of the individual changes or change direction; antagonistic, if the overall change is smaller than the sum of the individual changes. In contrast to Piggott et al., we here did not explicitly include the definition of positive and negative antagonism and synergism. The interaction analysis is based on numerical simulations with a threshold of 1%. Therefore, an effect is termed additive as long as the overall change is within a threshold of 1% around the sum. In the algorithm, it is first checked whether the additive case is true, followed by a check for synergy. If these are not true, the interaction is classified as antagonistic.

## Acknowledgments

M.S. was funded by a PhD fellowship of the graduate school “Computational Systems Biology” of the German Research Foundation (DFG Graduiertenkolleg 1772). The project was supported by a grant from the e:Med-program of the German Federal Ministry of Education and Research (BMBF): SYSMED-NB to J.W. (grant number: 01ZX1607F) and A.S (grant number:01ZX1607C). The manuscript is partly based on data generated in a project funded by German Cancer Aid (70113455, PI: A.S.). The funders had no role in study design, data collection and analysis, decision to publish, or preparation of the manuscript.

## Author contributions

J.W. and A.S designed the project. J.W., U.B., and K.B. supervised the analyses. Data analysis was performed by K.B. and M.S., quantitative dynamic modeling, and model analysis were performed by M.S., Visualization was done by M.S. and J.W., M.S. and J.W. wrote and all authors approved the manuscript.

