## Supplement for "A computational model elucidates the effect of oncogene induced expression alterations on the energy metabolism of neuroblastoma"

#### Ordinary differential equations

$$\frac{\mathrm{d}[glc]}{\mathrm{d}t} = \mathrm{v}_{\mathrm{glc\_tp}} - \mathrm{v}_{\mathrm{HK}}$$

$$\frac{\mathrm{d}[glc6p]}{\mathrm{d}t} = \mathrm{v}_{\mathrm{HK}} - \mathrm{v}_{\mathrm{GPI}}$$

$$\frac{\mathrm{d}[fru6p]}{\mathrm{d}t} = \mathrm{v}_{\mathrm{GPI}} - \mathrm{v}_{\mathrm{PFK1}}$$

$$\frac{\mathrm{d}[fru16bp]}{\mathrm{d}t} = \mathrm{v}_{\mathrm{PFK1}} - \mathrm{v}_{\mathrm{ALD}}$$

$$\frac{\mathrm{d}[dhap]}{\mathrm{d}t} = \mathrm{v}_{\mathrm{ALD}} - \mathrm{v}_{\mathrm{TPI}}$$

$$\frac{\mathrm{d}[gap]}{\mathrm{d}t} = \mathrm{v}_{\mathrm{ALD}} + \mathrm{v}_{\mathrm{TPI}} - \mathrm{v}_{\mathrm{GAPDH}}$$

$$\frac{\mathrm{d}[bpg]}{\mathrm{d}t} = \mathrm{v}_{\mathrm{GAPDH}} - \mathrm{v}_{\mathrm{PGK}}$$

$$\frac{\mathrm{d}[3pg]}{\mathrm{d}t} = \mathrm{v}_{\mathrm{PGK}} - \mathrm{v}_{\mathrm{PGM}}$$

$$\frac{\mathrm{d}[2pg]}{\mathrm{d}t} = \mathrm{v}_{\mathrm{PGM}} - \mathrm{v}_{\mathrm{ENO}}$$

$$\frac{\mathrm{d}[pep]}{\mathrm{d}t} = \mathrm{v}_{\mathrm{ENO}} - \mathrm{v}_{\mathrm{PK}}$$

$$\frac{\mathrm{d}[pyr]}{\mathrm{d}t} = \mathrm{v}_{\mathrm{PK}} - \mathrm{v}_{\mathrm{LDH}} - \mathrm{v}_{\mathrm{PDH}}$$

$$\frac{\mathrm{d}[lac]}{\mathrm{d}t} = \mathrm{v}_{\mathrm{LDH}} - \mathrm{v}_{\mathrm{lac\_tp}}$$

$$\frac{\mathrm{d}[acoa]}{\mathrm{d}t} = \mathrm{v_{PDH}} - \mathrm{v_{TCA}}$$

$$\frac{\mathrm{d}[adp]}{\mathrm{d}t} = \mathrm{v_{HK}} + \mathrm{v_{PFK1}} - \mathrm{v_{PGK}} - \mathrm{v_{PK}} - \mathrm{v_{TCA}} - 14 \cdot \mathrm{v_{Resp}} + \mathrm{v_{ATP\_cons}}$$

$$\frac{\mathrm{d}[atp]}{\mathrm{d}t} = -\mathrm{v_{HK}} - \mathrm{v_{PFK1}} + \mathrm{v_{PGK}} + \mathrm{v_{PK}} + \mathrm{v_{TCA}} + 14 \cdot \mathrm{v_{Resp}} - \mathrm{v_{ATP\_cons}}$$

$$\frac{\mathrm{d}[nad]}{\mathrm{d}t} = -\mathrm{v_{GAPDH}} + \mathrm{v_{LDH}} - \mathrm{v_{PDH}} - 4 \cdot \mathrm{v_{TCA}} + 6 \cdot \mathrm{v_{Resp}}$$

$$\frac{\mathrm{d}[nadh]}{\mathrm{d}t} = \mathrm{v_{GAPDH}} - \mathrm{v_{LDH}} + \mathrm{v_{PDH}} + 4 \cdot \mathrm{v_{TCA}} - 6 \cdot \mathrm{v_{Resp}}$$

$$\frac{\mathrm{d}[p]}{\mathrm{d}t} = -\mathrm{v_{GAPDH}} - \mathrm{v_{TCA}} - 14 \cdot \mathrm{v_{Resp}} + \mathrm{v_{ATP\_cons}}$$

$$\frac{\mathrm{d}[o2]}{\mathrm{d}t} = -3 \cdot \mathrm{v_{Resp}} + \mathrm{v_{o2\_tp}}$$

#### Conservation relations

The model includes three conservation relations. These include those for ATP and ADP as well as NAD and NADH since these are only converted into each other but not de novo synthesized in this model. The third one represents a conservation of the total phosphate availability and includes phosphate, ATP and all glycolytic intermediates from Glc6P to PEP.

| **Name** | **Included metabolites** |
|---|---|
| Total adenosine phosphates | atp+adp |
| Total redox metabolites | nad+nadh |
| Total phosphate | atp+p+glc6p+fru6p+2*fru16bp+gap+dhap+2*bpg+3pg+2pg+pep |

#### Rate equations

Most reactions are described by reversible Michaelis-Menten type kinetics. Oxygen transport and ATP consumption are described by mass-action kinetics. The mitochondrial reactions (PDH, TCA cycle, respiration) are modeled as irreversible. The oxygen transport is described by mass action kinetics, since it is based on diffusion. Since the complex GAPDH reaction was already modeled in a reduced form in Marín-Hernández et al. (2011), we here use the same rate law. The lumped reactions for the TCA cycle and respiration are adapted from Shestov et al. (2014). The $k_m$,$k_i$, $k_a$ and $k_{eq}$ values are specific for each reaction. Additional subscripts indicating the reaction name were omitted in the rate laws for easier readability.
ATP and ADP are representing also other nucleotide phosphates (e.g. GTP and GDP) and NAD and NADH are representing all redox metabolites (e.g. FAD and FADH2).

**Glucose transporter**

$$v_{glc\_tp} = \frac{\frac{vmf_{glc\_tp}}{k_m} \cdot ([glc\_ext] - \frac{[glc]}{k_{eq}})}{1 + \frac{[glc\_ext]}{k_m} + \frac{[glc]}{k_m}}$$

**Hexokinase**

$$v_{HK} = \frac{\frac{vmf_{HK}}{k_{m,glc} \cdot k_{m,atp}} \cdot ([glc] \cdot [atp] - \frac{[glc6p] \cdot [adp]}{k_{eq}})}{1 + \frac{[glc]}{k_{m,glc}} + \frac{[atp]}{k_{m,atp}} + \frac{[glc] \cdot [atp]}{k_{m,glc} \cdot k_{m,atp}} + \frac{[glc6p]}{k_{m,glc6p}} + \frac{[adp]}{k_{m,adp}} + \frac{[glc6p] \cdot [adp]}{k_{m,glc6p} \cdot k_{m,adp}}} \cdot \frac{k_{i,glc6p}}{k_{i,glc6p} + [glc6p]}$$

**Glucose phosphate isomerase**

$$v_{GPI} = \frac{\frac{vmf_{GPI}}{k_{m,glc6p}} \cdot ([glc6p] - \frac{[fru6p]}{k_{eq}})}{1 + \frac{[glc6p]}{k_{m,glc6p}} + \frac{[fru6p]}{k_{m,fru6p}}}$$

**Phosphofructokinase**

$$v_{PFK1} = \frac{\frac{vmf_{PFK1}}{k_{m,fru6p} \cdot k_{m,atp}} \cdot ([fru6p] \cdot [atp] - \frac{[fru16bp] \cdot [adp]}{k_{eq}})}{1 + \frac{[fru6p]}{k_{m,fru6p}} + \frac{[atp]}{k_{m,atp}} + \frac{[fru6p] \cdot [atp]}{k_{m,fru6p} \cdot k_{m,atp}} + \frac{[fru16bp]}{k_{m,fru16bp}} + \frac{[adp]}{k_{m,adp}} + \frac{[fru16bp] \cdot [adp]}{k_{m,fru16bp} \cdot k_{m,adp}}} \cdot \frac{k_{i,atp}}{k_{i,atp} + [atp]}$$

**Aldolase**

$$v_{ALD} = \frac{\frac{vmf_{ALD}}{k_{m,fru16bp}} \cdot ([fru16p] - \frac{[dhap] \cdot [gap]}{k_{eq}})}{1 + \frac{[fru16bp]}{k_{m,fru16bp}} + \frac{[dhap]}{k_{m,dhap}} + \frac{[gap]}{k_{m,gap}} + \frac{[dhap] \cdot [gap]}{k_{m,dhap} \cdot k_{m,gap}}}$$

**Triose-phosphate isomerase**

$$v_{TPI} = \frac{\frac{vmf_{TPI}}{k_{m,dhap}} \cdot ([dhap] - \frac{[gap]}{k_{eq}})}{1 + \frac{[dhap]}{k_{m,dhap}} + \frac{[gap]}{k_{m,gap}}}$$

**Glyceraldehyde 3-phosphate dehydrogenase**

$$v_{GAPDH} = \frac{\frac{vmf_{GAPDH}}{k_{m,nad} \cdot k_{m,gap} \cdot k_{m,p}} \cdot ([nad] \cdot [gap] \cdot [p] - \frac{[nadh] \cdot [bpg]}{k_{eq}})}{1 + \frac{[nad]}{k_{m,nad}} + \frac{[nad] \cdot [gap]}{k_{m,nad} \cdot k_{m,gap}} + \frac{[nad] \cdot [gap] \cdot [p]}{k_{m,nad} \cdot k_{m,gap} \cdot k_{m,p}} + \frac{[nadh]}{k_{m,nadh}} + \frac{[nadh] \cdot [bpg]}{k_{m,nadh} \cdot k_{m,bpg}}}$$

##### Phosphoglycerate kinase

$$v_{PGK} = \frac{\frac{vmf_{PGK}}{k_{m,bpg} \cdot k_{m,adp}} \cdot ([bpg] \cdot [adp] - \frac{[3pg] \cdot [atp]}{k_{eq}})}{1 + \frac{[bpg]}{k_{m,bpg}} + \frac{[adp]}{k_{m,adp}} + \frac{[bpg] \cdot [adp]}{k_{m,bpg} \cdot k_{m,adp}} + \frac{[3pg]}{k_{m,3pg}} + \frac{[atp]}{k_{m,atp}} + \frac{[3pg] \cdot [atp]}{k_{m,3pg} \cdot k_{m,atp}}}$$

##### Phosphoglycerate mutase

$$v_{PGM} = \frac{\frac{vmf_{PGM}}{k_{m,3pg}} \cdot ([3pg] - \frac{[2pg]}{k_{eq}})}{1 + \frac{[3pg]}{k_{m,3pg}} + \frac{[2pg]}{k_{m,2pg}}}$$

##### Enolase

$$v_{ENO} = \frac{\frac{vmf_{ENO}}{k_{m,2pg}} \cdot ([2pg] - \frac{[pep]}{k_{eq}})}{1 + \frac{[2pg]}{k_{m,2pg}} + \frac{[pep]}{k_{m,pep}}}$$

##### Pyruvate kinase

$$v_{PK} = \frac{\frac{vmf_{PK}}{k_{m,pep} \cdot k_{m,adp}} \cdot ([pep] \cdot [adp] - \frac{[pyr] \cdot [atp]}{k_{eq}})}{1 + \frac{[pep]}{k_{m,pep}} + \frac{[adp]}{k_{m,adp}} + \frac{[pep] \cdot [adp]}{k_{m,pep} \cdot k_{m,adp}} + \frac{[pyr]}{k_{m,pyr}} + \frac{[atp]}{k_{m,atp}} + \frac{[pyr] \cdot [atp]}{k_{m,pyr} \cdot k_{m,atp}}} \cdot (1 + \frac{[fru16bp]}{k_{a,fru16p}})$$

##### Lactate dehydrogenase

$$v_{LDH} = \frac{\frac{vmf_{LDH}}{k_{m,pyr} \cdot k_{m,nadh}} \cdot ([pyr] \cdot [nadh] - \frac{[lac] \cdot [nad]}{k_{eq}})}{1 + \frac{[pyr]}{k_{m,pyr}} + \frac{[nadh]}{k_{m,nadh}} + \frac{[pyr] \cdot [nadh]}{k_{m,pyr} \cdot k_{m,nadh}} + \frac{[lac]}{k_{m,lac}} + \frac{[nad]}{k_{m,nad}} + \frac{[lac] \cdot [nad]}{k_{m,lac} \cdot k_{m,nad}}}$$

##### Lactate transporter

$$v_{lac\_tp} = \frac{\frac{vmf_{lac\_tp}}{k_m} \cdot ([lac] - \frac{[lac\_ext]}{k_{eq}})}{1 + \frac{[lac]}{k_m} + \frac{[lac\_ext]}{k_m}}$$

##### Pyruvate dehydrogenase

$$v_{PDH} = \frac{vmax_{PDH} \cdot [pyr] \cdot [nad]}{k_{m,pyr} \cdot k_{m,nad} + [pyr] \cdot k_{m,nad} + [nad] \cdot k_{m,pyr} + [pyr] \cdot [nad]}$$

**TCA cycle** $$\mathrm{v_{TCA}} = \frac{\mathrm{vmax_{CC}} \cdot \frac{[\mathrm{acoa}] \cdot [\mathrm{nad}] \cdot [\mathrm{adp}] \cdot [\mathrm{p}]}{\mathrm{k_{m,acoa}} \cdot \mathrm{k_{m,nad}} \cdot \mathrm{k_{m,adp}} \cdot \mathrm{k_{m,p}}}}{(1 + \frac{[\mathrm{acoa}]}{\mathrm{k_{m,acoa}}}) \cdot (1 + \frac{[\mathrm{nad}]}{\mathrm{k_{m,nad}}}) \cdot (1 + \frac{[\mathrm{adp}]}{\mathrm{k_{m,adp}}}) \cdot (1 + \frac{[\mathrm{p}]}{\mathrm{k_{m,p}}})}$$

**Respiration**

$$\mathrm{v_{Resp}} = \frac{\mathrm{vmax_{Resp}} \cdot \frac{[\mathrm{o2}] \cdot [\mathrm{nadh}] \cdot [\mathrm{adp}] \cdot [\mathrm{p}]}{\mathrm{k_{m,o2}} \cdot \mathrm{k_{m,nadh}} \cdot \mathrm{k_{m,adp}} \cdot \mathrm{k_{m,p}}}}{(1 + \frac{[\mathrm{o2}]}{\mathrm{k_{m,o2}}}) \cdot (1 + \frac{[\mathrm{nadh}]}{\mathrm{k_{m,nadh}}}) \cdot (1 + \frac{[\mathrm{adp}]}{\mathrm{k_{m,adp}}}) \cdot (1 + \frac{[\mathrm{p}]}{\mathrm{k_{m,p}}})}$$

**$O_2$ transport**

$$\mathrm{v_{o2\_tp}} = \mathrm{k1_{o2\_tp}} \cdot ([\mathrm{o2_{ext}}] - [\mathrm{o2}])$$

**ATP consumption**

$$\mathrm{v_{ATP\_cons}} = \mathrm{k1_{atp\_cons}} \cdot [\mathrm{atp}]$$

#### Parameters

The over 80 kinetic parameters including Michaelis-Menten constants and maximal velocities should be assigned in a neuroblastoma specific way. Where available experimentally determined values for neuroblastoma cells were used. km values were either estimated for neuroblastoma cells or are taken from the BRENDA (Jeske et al., 2019) and SABIO-RK (Wittig et al., 2012) databases. Here data for *Homo sapiens* were used whenever possible. Equilibrium constants are taken from literature (Holzhütter, 2004). The initial range of these values was set based on typical values for mammalian cells (as given in Mulukutla et al., 2014; Roy and Finley, 2017; Shestov et al., 2014), The maximal velocities, which are commonly assumed to be cell type specific, were then fitted using metabolomic data (for glc, g6p, fru16bp, dhap, 3pg, pep, pyr, lac, acoa, adp, nad, nadh) and the extracellular flux data (ECAR and OCR values from three independent experiments), each for three different glucose concentrations in the medium (5.55, 11.11 and 25 mM), from a neuroblastoma cell line as described in (Tjaden et al., 2020) (see Methods).

Table S.1: **List of parameters of the reference model.** The parameters are sorted by type and then alphabetically. The sources are: I for values from Holzhütter, 2004, II for parameters experimentally determined for this cell line (Tjaden et al., 2020), III for fitted parameters, IV for literature/database derived parameters (Jeske et al., 2019, Wittig et al., 2012). All concentrations, Michaelis Menten constants, inhibition and activation constants are given in mM, maximal velocities in mM/min. The units of the equilibrium constants depend on the number of products and substrates, for same number of substrates and products there is no unit. The table continues on the next page.

| Parameter | Value | Source |
|---|---|---|
| keq_ALD | 0.114 | I |
| keq_ENO | 1.7 | I |
| keq_GAPDH | 1.92E-04 | I |
| keq_glctp | 1 | I |
| keq_GPI | 0.3922 | I |
| keq_HK | 3.90E+03 | I |
| keq_lactp | 1 | I |
| keq_LDH | 9.09E+03 | I |
| keq_PFK1 | 1.00E+05 | I |
| keq_PGK | 1460 | I |
| keq_PGM | 0.1449 | I |
| keq_PK | 1.38E+04 | I |
| keq_TPI | 4.07E-02 | I |
| glc_ext | 11.11 | II |
| lac_ext | 0.5 | I |
| o2_ext | 0.181 | I |

| Parameter | Value | Source |
|---|---|---|
| k1_atp_cons | 1.11 | III |
| k1_o2tp | 10.28 | III |
| vmax_CC | 3.79 | III |
| vmax_PDH | 1.89 | III |
| vmax_resp | 3.48 | III |
| vmf_ALD | 5474.59 | III |
| vmf_ENO | 5906.76 | III |
| vmf_GAPDH | 10000.00 | III |
| vmf_GPI | 9998.71 | III |
| vmf_HK | 4.89 | III |
| vmf_LDH | 9999.81 | III |
| vmf_PFK1 | 4.98 | III |
| vmf_PGK | 9999.41 | III |
| vmf_PGM | 8057.65 | III |
| vmf_PK | 23.00 | III |
| vmf_TPI | 8665.75 | III |
| vmf_glctp | 1.79 | III |
| vmf_lactp | 10000.00 | III |

| Parameter | Value | Source |
|---|---|---|
| km_ald_dhap | 0.095 | IV |
| km_ald_f16bp | 0.0107 | IV |
| km_ald_gap | 1.17 | IV |
| km_cc_acoa | 0.278 | III |
| km_cc_adp | 3.599 | III |
| km_cc_nad | 0.071 | III |
| km_cc_p_i | 0.289 | III |
| km_eno_2pg | 0.03 | IV |
| km_eno_pep | 0.702 | IV |
| km_gapdh_bpg | 0.01 | IV |
| km_gapdh_gap | 0.149 | IV |
| km_gapdh_nad | 0.071 | IV |
| km_gapdh_nadh | 0.01 | IV |
| km_gapdh_p | 4 | IV |
| Km_glctp | 1.5 | IV |
| km_gpi_f6p | 0.0635 | IV |
| km_gpi_g6p | 0.445 | IV |
| km_hk_adp | 11.1 | IV |
| km_hk_atp | 0.46 | II |
| km_hk_g6p | 2 | IV |
| km_hk_glc | 0.064 | II |
| ki_hk_g6p | 0.24 | IV |
| km_lactp | 4.68 | IV |
| km_ldh_lac | 0.94 | IV |
| km_ldh_nad | 0.311 | IV |
| km_ldh_nadh | 0.014 | IV |
| km_ldh_pyr | 0.08 | II |
| km_pdh_nad | 0.033 | IV |
| km_pdh_pyr | 0.03 | IV |
| km_pfk_adp | 0.49 | IV |
| km_pfk_atp | 0.05 | IV |
| km_pfk_f16bp | 16.7 | IV |
| km_pfk_f6p | 0.45 | IV |
| ki_pfk_atp | 1.6 | IV |

| Parameter | Value | Source |
|---|---|---|
| km_pgk_3pg | 0.009 | IV |
| km_pgk_adp | 0.2 | IV |
| km_pgk_atp | 0.07 | IV |
| km_pgk_bpg | 0.024 | IV |
| km_pgm_2pg | 0.041 | IV |
| km_pgm_3pg | 0.4 | IV |
| km_pk_adp | 0.17 | IV |
| km_pk_atp | 0.35 | IV |
| km_pk_pep | 1.1 | IV |
| km_pk_pyr | 0.48 | IV |
| ka_pk_f16bp | 0.1 | IV |
| km_resp_adp | 0.001 | III |
| km_resp_nadh | 0.001 | III |
| km_resp_o2 | 0.001 | III |
| km_resp_p_i | 10 | III |
| km_tpi_dhap | 0.17 | IV |
| km_tpi_gap | 1.76 | II |

### Supplemental Figures

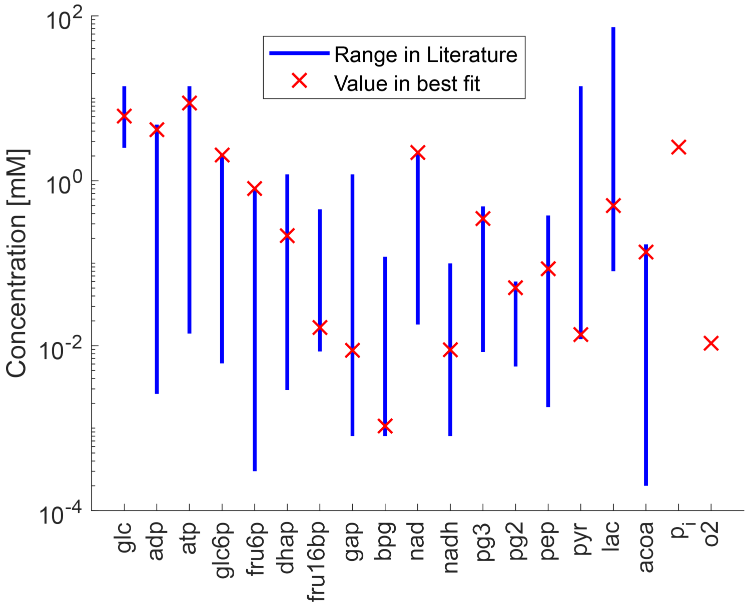

Figure S.1: **Comparison of fitted model concentrations to known literature values.** Experimental range, shown by the blue bars, are taken from Roy and Finley (2017), these represent concentrations in different mammalian cell lines, mostly cancer cell lines. Therefore, they are not representative for a single cell type or neuroblastoma. Oxygen (o2) and phosphate ($p_i$) are not included in this experimental dataset. The red crosses represent the steady state concentrations of the model with the best parameter fit. The steady state concentrations of the best fit are within the defined range from literature.

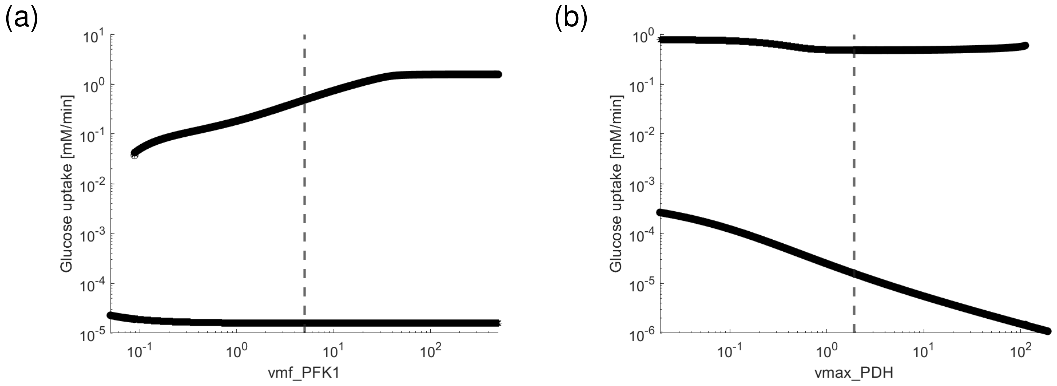

Figure S.2: **Coexistence of two stable steady states for additional parameters.** Stable steady state values in dependence of one parameter, for (a) maximal velocity of the PFK reaction and (b) the maximal velocity of the PDH reaction. The dashed line indicates the fitted value of the respective parameter. All other parameters are at their reference values (Supplemental Table S.1).

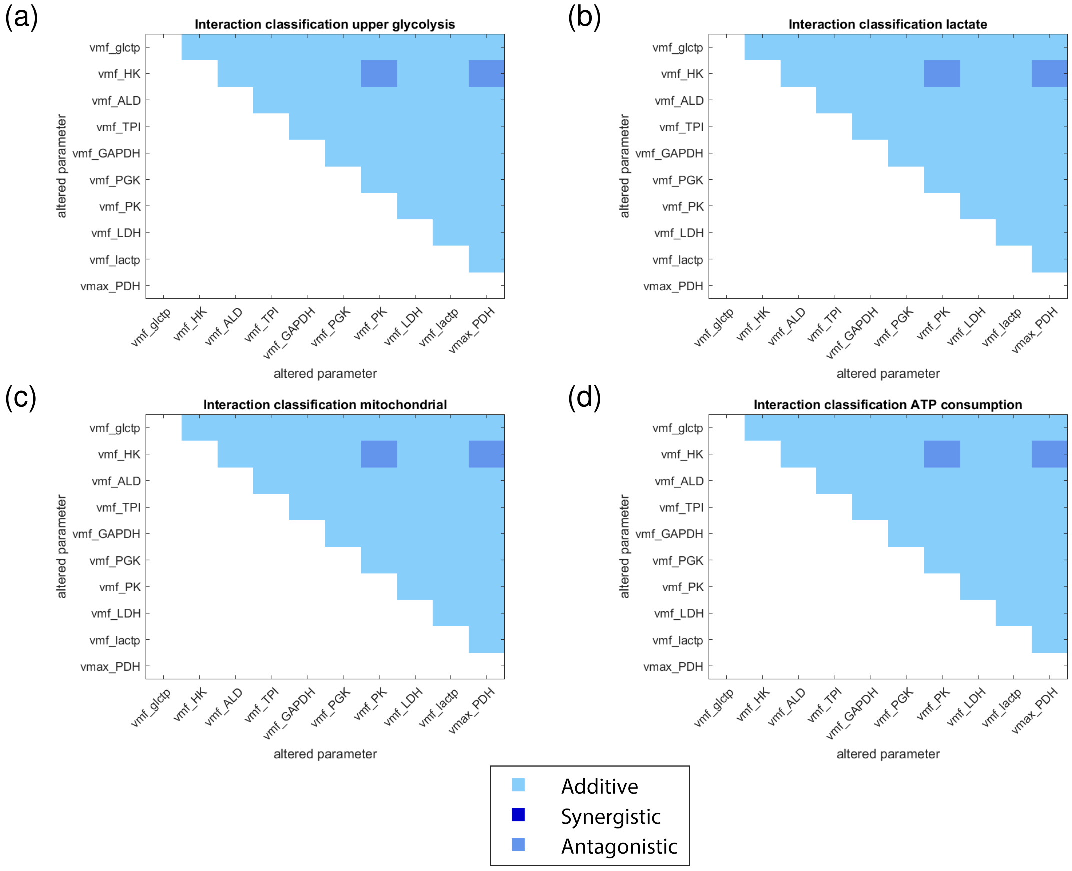

Figure S.3: **Interaction analysis of MYCN targets.** All MYCN targets are altered by a MYCN factor of two and the interaction effect between each possible pair of MYCN targets is analyzed. These effects are classified as either additive (light blue), synergistic (dark blue) or antagonistic (intermediate blue). While most effects are additive, two antagonistic cases can also be observed.

#### Supplemental Tables

Table S.2: **Steady state concentrations for the reference and low flux steady state.** All concentrations are given in mM.

| Metabolite | Steady state concentration [mM] Reference state | Low flux state |
|---|---|---|
| glc | 6.082 | 11.110 |
| glc6p | 2.052 | 2.134 |
| fru6p | 0.805 | 0.837 |
| fru16bp | 0.017 | 4.198 |
| dhap | 0.216 | 3.429 |
| gap | 0.009 | 0.140 |
| bpg | 0.001 | 7.70E-11 |
| 3pg | 0.349 | 1.47E-07 |
| 2pg | 0.050 | 2.11E-08 |
| pep | 0.086 | 3.56E-08 |
| pyr | 0.014 | 1.23E-06 |
| lac | 0.500 | 0.500 |
| acoa | 0.137 | 0.025 |
| o2 | 0.011 | 0.181 |
| atp | 8.776 | 0.001 |
| adp | 4.168 | 12.943 |
| nad | 2.202 | 0.048 |
| nadh | 0.009 | 2.163 |
| p | 2.559 | 1.33E-04 |

Table S.3: **Steady state fluxes for different MYCN factors.**

| | Steady state flux [mM/min] | | | |
|---|---|---|---|---|
| | Reference | MYCN factor | | |
| Flux | state | 1.1 | 1.5 | 2 |
| glc uptake | 0.48 | 0.50 | 0.56 | 0.62 |
| lac release | 0.38 | 0.42 | 0.57 | 0.75 |
| o2 uptake | 1.75 | 1.73 | 1.61 | 1.47 |
| ATP consumption | 9.71 | 9.62 | 9.17 | 8.57 |

Table S.4: **Steady state concentrations for different MYCN factors.**

| Metabolite | Steady state concentration [mM] Reference state | MYCN factor 1.5 | MYCN factor 2 |
|---|---|---|---|
| glc | 6.082 | 7.045 | 7.615 |
| glc6p | 2.052 | 2.736 | 3.309 |
| fru6p | 0.805 | 1.073 | 1.298 |
| fru16bp | 0.017 | 0.003 | 0.001 |
| dhap | 0.216 | 0.085 | 0.038 |
| gap | 0.009 | 0.003 | 0.002 |
| bpg | 0.001 | 0.001 | 0.0004 |
| 3pg | 0.349 | 0.282 | 0.222 |
| 2pg | 0.050 | 0.041 | 0.032 |
| pep | 0.086 | 0.069 | 0.055 |
| pyr | 0.014 | 0.023 | 0.033 |
| lac | 0.500 | 0.500 | 0.500 |
| acoa | 0.137 | 0.114 | 0.095 |
| o2 | 0.011 | 0.024 | 0.038 |
| atp | 8.776 | 8.285 | 7.740 |
| adp | 4.168 | 4.659 | 5.204 |
| nad | 2.202 | 2.206 | 2.208 |
| nadh | 0.009 | 0.005 | 0.004 |
| p | 2.559 | 2.358 | 2.239 |

Table S.5: **Leave-one-out-analysis for MYCN targets.** All MYCN targets except one are altered by a factor of 2. For comparison the top row shows the steady state fluxes for the case that all targets are altered, the last row shows the steady state if only five (glctp, HK, PGK, PK and PDH) of the targets are altered. Targets that lead to notable changes in the steady state fluxes are marked in grey.

| Left out | glc_tp [mM/min] | lac_tp [mM/min] | o2_tp [mM/min] | ATP_cons [mM/min] |
|---|---|---|---|---|
| None | 0.62 | 0.75 | 1.47 | 8.57 |
| vmf_glctp | 0.62 | 0.75 | 1.47 | 8.58 |
| vmf_HK | 0.49 | 0.40 | 1.71 | 9.52 |
| vmf_ALD | 0.62 | 0.75 | 1.47 | 8.57 |
| vmf_TPI | 0.62 | 0.75 | 1.47 | 8.57 |
| vmf_GAPDH | 0.62 | 0.75 | 1.47 | 8.57 |
| vmf_PGK | 0.62 | 0.75 | 1.46 | 8.56 |
| vmf_PK | 0.63 | 0.78 | 1.43 | 8.38 |
| vmf_LDH | 0.62 | 0.75 | 1.47 | 8.57 |
| vmf_lactp | 0.62 | 0.75 | 1.47 | 8.57 |
| vmax_PDH | 0.61 | 0.72 | 1.50 | 8.74 |
| Simultaneous leave-out of 5 targets | 0.62 | 0.75 | 1.47 | 8.57 |
